# A sensitized model of thrombosis validates known multigenic relationships and suggests novel modifiers of hemostasis

**DOI:** 10.64898/2026.05.11.724445

**Authors:** Steven J. Grzegorski, Yang Liu, Catherine E. Richter, Murat Yaman, Andy H. Vo, Xinge Yu, Anna R. Dahlgren, Hasam Madarati, Alexander P. Friedmann, Ida Surakka, Paul Y. Kim, Colin A. Kretz, Jordan A. Shavit

## Abstract

**Background:** Venous thromboembolism is a major cause of morbidity and mortality. Despite identification of risk factors, not all individuals with thrombophilia develop thrombosis. Understanding the multigenic factors modifying this incomplete penetrance would help guide patient care.

**Methods:** The zebrafish has a conserved hemostatic system and is amenable to large genetic studies. Loss of antithrombin III (At3) in zebrafish leads to an early consumptive coagulopathy and lethality in adulthood. Using this genetic background as a sensitized model we performed a dominant unbiased genome-wide N-ethyl-N-nitrosourea (ENU) mutagenesis screen followed by whole genome sequencing (WGS). We used survival studies, laser-mediated endothelial injury, and *ex vivo* protein assays to validate hits.

**Results:** ENU-treated *at3*^+/−^ males were crossed with *at3*^+/−^ females to produce 4,030 total offspring (1.5x genome coverage). Four permanent lines transmitting a survival benefit beyond 7 months were identified and sequenced. A candidate screen of 63 known coagulation-related loci revealed a missense mutation, C504F, in a highly conserved residue of the prothrombin (*F2*) heavy chain, which was validated through genetic and biochemical studies. Evaluation of UK Biobank electronic health record (EHR) data was underpowered to detect interactions between *F2* and *AT3* due to minmal deleterious mutations. Mutations produced through genome editing revealed that heterozygosity for factor X and plasminogen also modified *at3*^−/−^, resulting in reduced lethality. The three remaining lines had no coagulation-related variants segregating with survival, suggesting the presence of novel modifier loci.

**Conclusions:** Unbiased genome-wide screening identified a modifier of thrombosis. This demonstrated that re-balancing of hemostasis to mitigate thrombosis is conserved in zebrafish, including an unexpected role for fibrinolysis. This interaction was not detected even in a large human dataset, establishing the continued benefit of the zebrafish model. Finally, we found evidence for novel loci outside of the canonical coagulation cascade that may be new targets for diagnosis or treatment.

## Introduction

Venous thromboembolism (VTE) is defined by the formation of pathologic blood clots in deep veins with the risk of embolization to the pulmonary vasculature that is estimated to cause 60,000-100,000 American deaths per year^1,2^. Antithrombin III (AT3) is the primary endogenous inhibitor of coagulation, but due to unknown genetic modifiers, AT3 deficiency is incompletely penetrant, with estimates of a 50-80% lifetime risk of VTE^3^.

The pathology of VTEs centers around disruption of the normal processes underlying hemostasis including formation, stabilization, and dissolution of blood clots at sites of vascular injury. Under physiologic conditions, vascular injury results in the activation of coagulation factor X (FX) to form the serine protease FXa through the extrinsic or intrinsic pathways^4^. FXa then binds its cofactor factor Va (FVa) on negatively charged surface in the presence of calcium to form the prothrombinase complex, which converts the zymogen prothrombin (*F2* gene) to the protease thrombin. Thrombin is the final effector protease of coagulation that activates platelets and converts soluble fibrinogen to insoluble fibrin strands, which stabilizes thrombi to minimize blood loss at the site of injury. Subsequent activation of the fibrinolytic system facilitates the dissolution of these clots through proteolytic degradation of fibrin mediated by the enzyme plasmin. Plasmin is generated when the zymogen plasminogen (PLG) is activated by plasminogen activators such as tissue-type plasminogen activator or urokinase-type plasminogen activator. AT3 prevents initiation of this process by binding to thrombin and FX.

At each of the described steps, environmental and genetic risk factors contribute to risk of thrombosis. Genetically, increased procoagulant activity (*F2* 20210G>A, factor V Leiden (R506Q)) or decreased anticoagulant activity (AT3 deficiency, protein C deficiency, and protein S deficiency) can lead to ectopic and/or excess clot formation resulting in intravascular thrombus formation with the risk of subsequent embolization^4–7^. PLG deficiency in mice leads to the development of multiorgan fibrin deposits^8^, while deficiency in humans results in fibrin-rich pseudomembranes without increased thrombotic risk^9^. Clinically, catheter-directed thrombolysis through upstream activation of PLG leads to improved outcomes in some patient populations, but is associated with an increased risk of bleeding^10^. A recent study demonstrated that deficiency of thrombiactivatable fibrinolysis inhibitor (TAFI, proCPB2), led to decreased size of venous thrombi and increased lung embolic burden in a mouse model of VTE^11^. Taken together, these data suggest that the overall burden of VTE is influenced by both the coagulation and fibrinolytic cascades.

VTE occurs within the complicated milieu that is blood and vasculature, and is modified by dozens of known and likely unknown factors. Uncovering novel genetic risk factors for VTE continues to be challenging in human populations^12^, and can be complemented by high throughput forward genetic screens in animal models.

Zebrafish are an established system for studying the basic science of hemostasis and thrombosis, demonstrating a high degree of conservation of the coagulation cascade^13–16^. We have previously shown a conserved physiologic role of AT3 in zebrafish^17^. Complete deficiency leads to a consumptive coagulopathy with spontaneous venous thrombosis in larvae, and early mortality due to intracardiac thrombi in adults. This clot burden results in 90% lethality by 7 months of age. With a strong history of forward genetics and a rapidly developing toolkit of reverse genetic technology^18^, the zebrafish offers a unique *in vivo* model for interrogating and discovering complex modifiers of thrombosis within and beyond the canonical coagulation cascade.

Using the *at3*^−/−^ thrombophilic zebrafish line, we pursued a two-pronged approach to identifying genetic modifiers of thrombosis. We first utilized an unbiased classical N-ethyl-N-nitrosourea (ENU) mutagenesis protocol to isolate novel mutations and loci^19,20^, including a variant in *f2*, which we have named *frost*. Despite the well-known relationship between AT3 and thrombin, a large human population-based dataset (UK Biobank) was still unable to detect the interaction discovered in this animal model. This was followed by a targeted genome editing approach to knock out known coagulation factors that would enhance or diminish survival. In the process we improved our understanding of thrombin biology, uncovered expected and unexpected risk mitigators not detectable in a human EHR, and identified candidate suppressor loci independent of known coagulation factors.

## Methods

### Animal Care

Zebrafish were maintained according to protocols approved by the University of Michigan Animal Care and Use Committee. All wild-type fish were a hybrid line generated by crossing AB and TL fish acquired from the Zebrafish International Resource Center. A hybrid line was selected to minimize strain specific effects and for improved reproductive success^21^. Tris-buffered tricaine methaneosulfate (Western Chemical) was used for anesthesia during all live animal experimental manipulation and for euthanasia. Knockout backgrounds of *at3, f2, f10* (FX), and *plg* have been previously described.^17,22–24^

### Genome-wide mutagenesis through N-ethyl-N-nitrosourea (ENU) treatment

Fish were maintained and chemically treated in accordance with a protocol approved by the University of Michigan Animal Care and Use Committee and similar to previously published methods^19,20^. Briefly, ENU was resuspended in 10 mM acetic acid to a concentration of 85.5 mM and diluted in 10 mM sodium phosphate buffer to a final concertation of 3.3 mM. Male fish were treated for one hour in 500 mL of solution. Fish were allowed to recover for one hour in phosphate buffer and then transferred to fish water with tricaine for another hour followed by a final 3-hour wash in fish water and tricaine. Fish were then transferred to clean fish water and left undisturbed overnight. This procedure was repeated for three cycles with one week of rest between periods with breeding beginning one week after the final treatment.

### Genotyping of mutant offspring

Whole larvae or adult fin biopsies were lysed in buffer (10 mM Tris-HCl, pH 8.0, 2 mM EDTA, 2% Triton X-100, 100 µg/mL proteinase K) for 2 hours at 55°C followed by incubation at 95°C for 5 minutes. One microliter of the resulting solution was used as template for gene specific PCR (Table 1) and analyzed by gel electrophoresis. For *frost* genotyping, the amplified product was digested for 1 hour with ApaLI prior to electrophoresis. Additionally, an rhAmp Assay (IDT) was designed to rapidly genotype for the *frost* prothrombin variant (Design ID: CD.GT.RQFP5275.2).

**Table 1.**
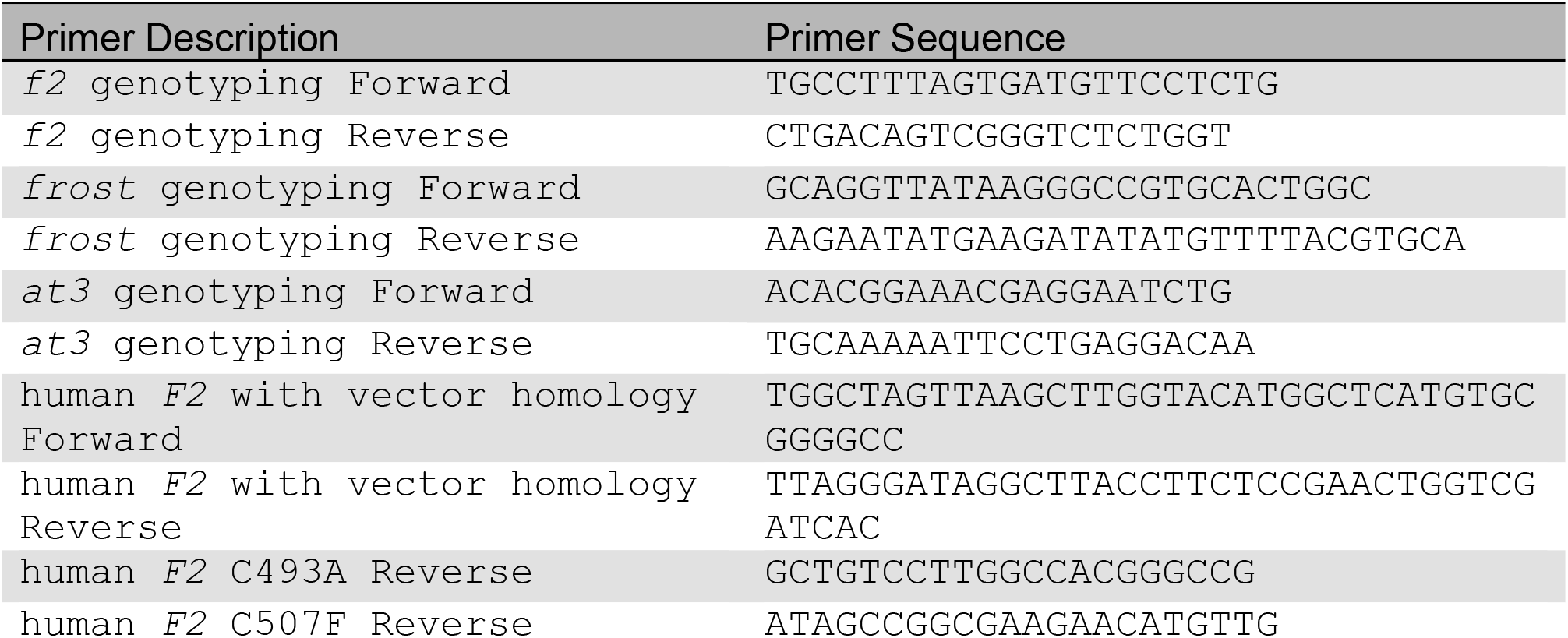
Oligonucleotide sequences.

### Whole genome sequencing (WGS) and variant calling

Tail DNA was extracted from adult zebrafish and purified using the Qiagen DNeasy Kit. The purified DNA was sent to Novogene for library preparation and sequencing on the Illumina short read platform. Whole genome data were aligned to the GRCz11 reference sequence^25^. Standard packages were selected to be consistent with the Broad Institute’s Genomic Analysis Toolkit (GATK) Best Practices^26^. Alignment was done using the Burrow-Wheelers aligner (bwa v0.7.15; bwa-mem algorithm)^27^ and converted to the bam file format using samtools (v1.9; view tool). Optical duplicates were identified, marked and sorted using the GATK (v4.1.0.0) tool. Notably, MarkDuplicateSparks is a Spark implementation for multi-core parallel processing with processing time improvements that scale linearly to 16 local cores. For this reason, major portions of this pipeline were optimized to run on 16 cores. Base qualities were recalibrated using the GATK tools BQSR and ApplyBQSR. Variants were called using a two-phase approach with the GATK tools HaplotypeCaller and GenotypeGVCFs. Additional annotation and post processing was done using SnpSift and SnpEff^28^. All large data processing was performed on the University of Michigan Great Lakes High Performance Computing Cluster.

### Laser-induced endothelial injury

Laser injury was performed as previously described^29^. Briefly, larvae were mounted in 0.8% low melt agarose and placed on an Olympus IX73 with attached Andor Technology Micropoint Pulsed Laser System (Belfast, United Kingdom). At 3 days post fertilization (dpf), 99 pulses were administered to the ventral endothelial surface of the posterior cardinal vein, 5 somites caudal to the anal pore. Two minutes after injury, larvae were examined for formation of an occlusive thrombus and scored as either occluded or not occluded.

### Human prothrombin mutant vector construction and protein expression

pcDNA3.1 containing the human *F2* cDNA from our previous study^24^ was modified using site-directed mutagenesis^30^ to generate the *frost* mutation at the homologous cysteine, C507F. As C507 forms a disulfide bond with C493, a corresponding set of vectors (C493A and C493A/C507F) were generated to control for the effect of a free cysteine (Table 1). Vectors were then expressed in HEK293T cells and the protein purified as previously described^24^.

### Prothrombin activation and activity

Assays on prothrombin and the *frost* variant were performed as previously described^24^. Purified prothrombin was activated using FXa and quantified by formation of DAPA-thrombin complexes. Thrombin activity was assayed using the substrates S-2238 and fibrinogen to check for active site maturation and macromolecular substrate recognition, respectively. Note that these studies were performed in parallel with a previously published analysis of a TALEN-generated *f2* mutation (*f2*^*Δ*^)^24^. Therefore, the control data for *frost* are the same as in the analysis of *f2*^*Δ*^.

### UK Biobank data analysis

For each individual in the UK Biobank (UKBB) with Haplotype Reference Consortium (HRC) imputed genotype data available (N=470,751), the VTE case status was derived using specific ICD10 codes: I80.1 or I80.2 (phlebitis and thrombophlebitis of the femoral vein or other deep vessels of lower extremities), I82.2 (embolism and thrombosis of vena cava), I26.0 or I26.9 (pulmonary embolism with or without acute cor pulmonale) and procedure codes L79.1 or L90.2 (open venous thrombectomy of lower limb vein or inferior vena cava filter insertion). This resulted in a total of 7,992 VTE cases and 462,759 controls. To identify individuals carrying deleterious mutations in the *F2, F10* and *AT3* genes, we applied WGS Annotation (WGSA) to all variants within these genes. Variants were classified as deleterious if their CADD (Combined Annotation Dependent Depletion) score exceeded the Mutation Significance Cutoff (MSC) at the 95% confidence interval (CI) as defined by the Human Gene Mutation Database (HGMD).

### Statistical Analysis

The occlusion data were analyzed using Mann-Whitney *U* or two-tailed Student’s *t* tests. Survival was evaluated by log-rank (Mantel-Cox) testing. Significance testing, graphs, and survival curves were produced with GraphPad Prism (v10.4.1; GraphPad Software, La Jolla, California). For the UKBB data, significance of the association between VTE prevalence and deleterious mutation carriers was assessed using Fisher’s exact test. This test was performed using the fisher.test() function in R (v4.3.3). Statistical significance was defined as p<0.05. All data were collected by observers blinded to condition or genotype.

## Results

### *frost*, a novel prothrombin mutation, results in substitution of a highly conserved residue

In order to survey for novel modifier genes, unbiased genome-wide chemical mutagenesis was performed on *at3*^+/−^ males. Subsequent breeding to *at3*^+/−^ females yielded 4030 surviving offspring which were genotyped between 2-5 months of age. Of these the resulting genotypes were 1248 *at3*^*+/+*^, 2516 *at3*^+/−^, and 266 *at3*^−/−^. Assuming 30 coding mutations per spermatogonia and a Mendelian expectation of ~1250 *at3*^−/−^ offspring, the mutagenesis achieved 1.5x coverage of the genome. Power analysis indicates that we covered roughly 78% of mutable loci, while 95% coverage would require~10,000 offspring. 53 of the homozygotes survived to 7 months, 23 were able to produce offspring and 4 lines (*frost, pearl, starr*, and *quinn*) stably transmitting a survival benefit were established (Figure 1). We performed WGS on the 4 stable lines. Variants that were unique to each line (absent in control fish, dbSNP v150, SNPFisher, and other sequenced lines) and present in at least 10 members of that line were annotated. An initial candidate-based approach found 3 lines with mutations in known clotting or related genes (of 63 genes surveyed from the Thrombogenomics consortium^31^). Only one mutation segregated with survival. This encoded a cysteine to phenylalanine change at the highly conserved position 504 of prothrombin (C504F; polyphen-2^32^ = 1.000 (G>T; phyloP^33^ = 9.69), Figure 2) in the *frost* line. An initial pool of 47 *at3*^−/−^ fish greater than 2 years of age from the *frost* lineage were genotyped for this mutation and 45 found to be carriers. A separate group of fish were genotyped at 2 months and followed for survival. This group again demonstrated a survival advantage when carrying the C504F allele (represented by *f2*^*C/F*^, Figure 3A).

**Figure 1.**
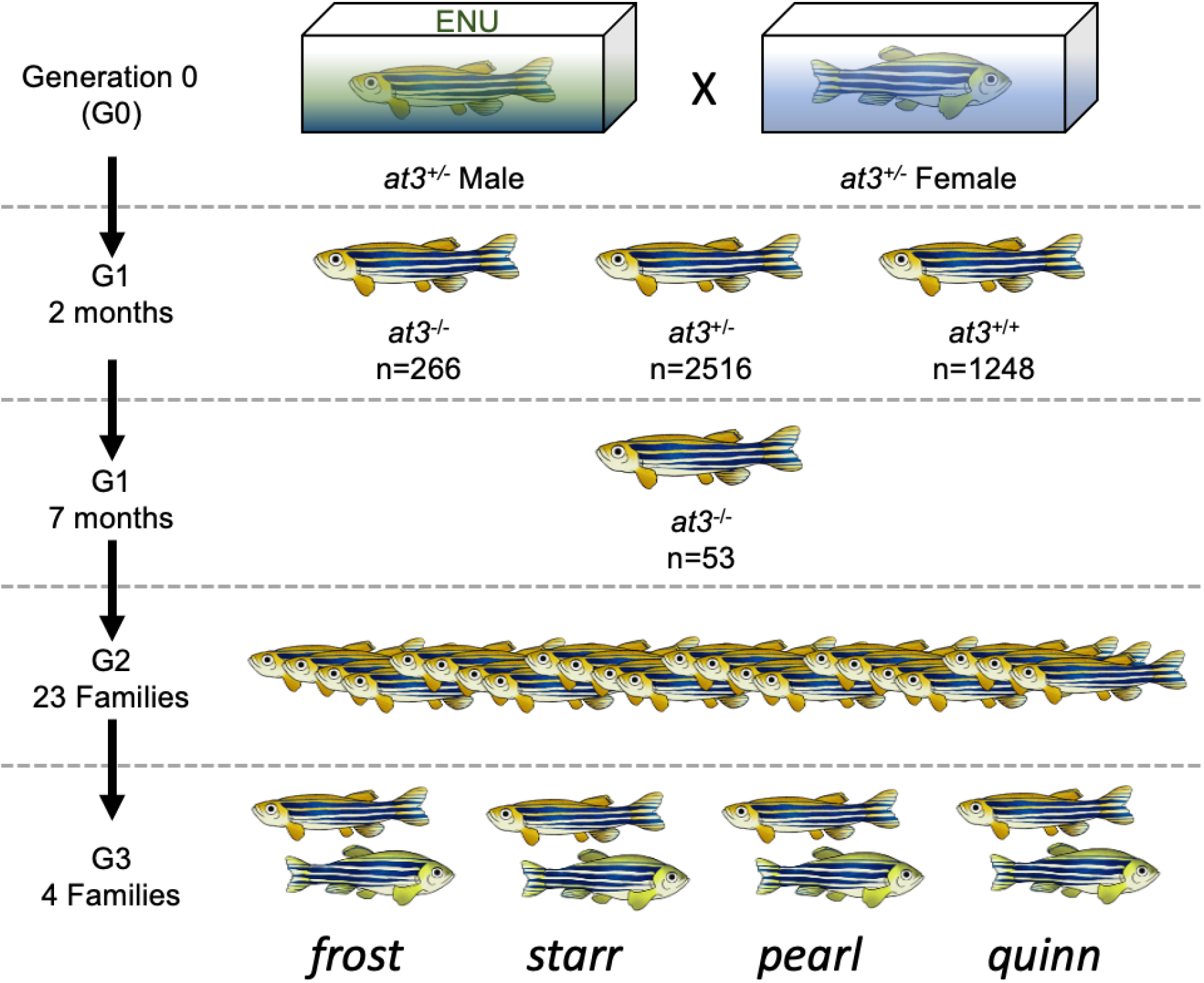
Overview of ENU mutagenesis. Mutagenized *at3*^+/−^ males were crossed to non-mutagenized females and 4030 offspring were produced with a predicted genomic coverage of 1.5x. At 2-5 months, *at3* deficient offspring were significantly underrepresented. By 7 months, 53 surviving fish remained and were selected for outcrossing. 23 fish produced viable offspring and from these 4 separate families were established with a transmittable survival benefit.

**Figure 2.**
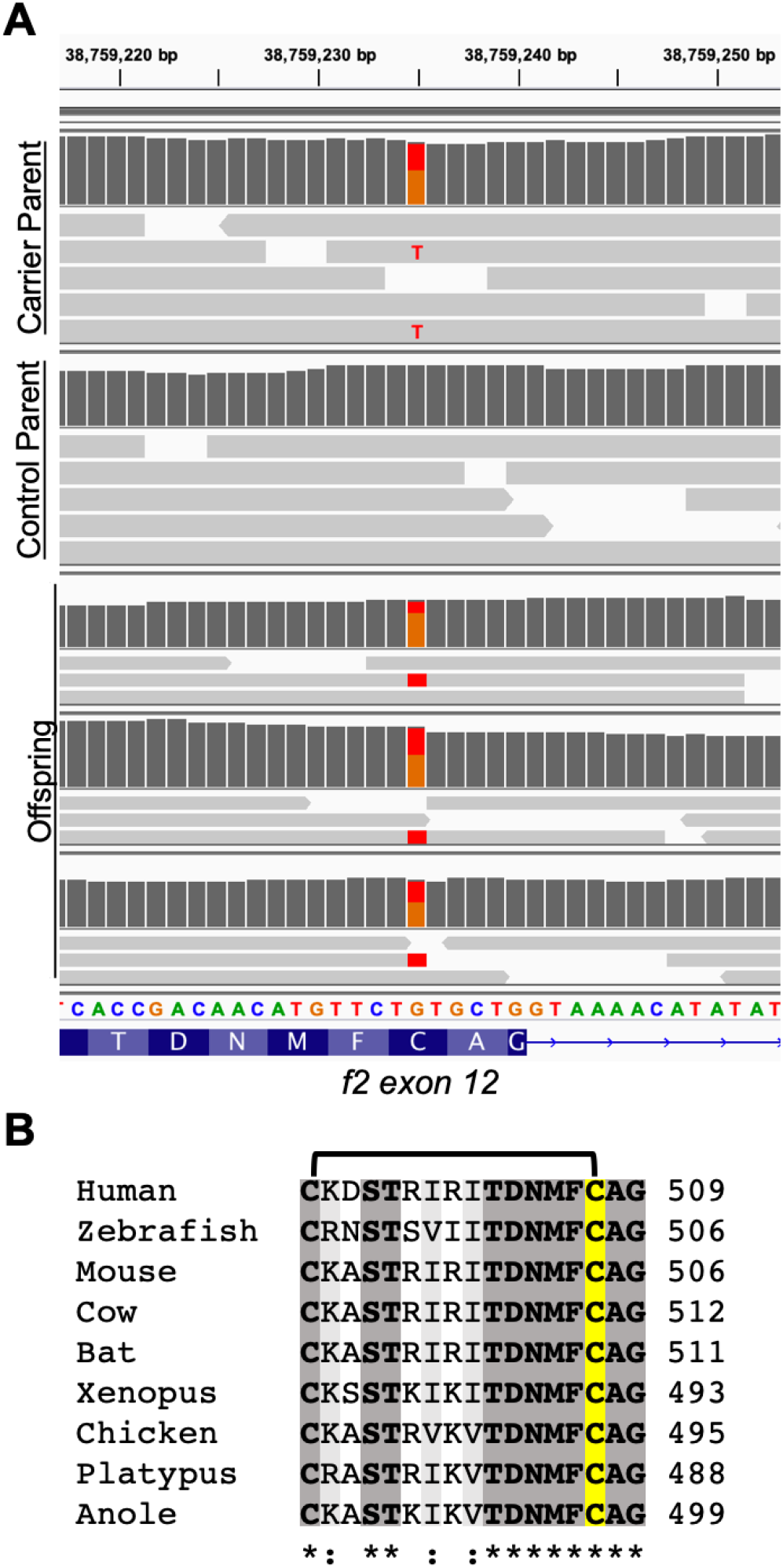
WGS identifies a missense mutation in the codon for a highly conserved cysteine of *f2*. (A) Integrative Genomics Viewer display of the short-read sequencing around the identified variant site in exon 12 of *f2*. Carrier and control parents and 3 surviving offspring were sequenced. Dark grey bars indicate coverage for each sample scaled to 35 reads. Light grey bars correspond to individual reads with single G>T mutations indicated by red labeling, and reference allele indicated in orange. (B) Amino acid sequence alignment of prothrombin residues surrounding the conserved cysteine in multiple organisms. Identity is shaded in dark grey and bolded. Homology is shaded in light grey. The cysteine of interest is highlighted in yellow and the disulfide bond annotated above the alignment.

### Confirmation that *frost* is a mutant allele of *f2*

Although these data are compelling, a formal possibility remained that another gene in linkage disequilibrium with *f2* could harbor the suppressor mutation. Therefore, to validate the C504F allele as the *frost* mutation, we mated our prothrombin knockout^24^ (referred to as *f2*^*Δ*^) into the non-mutagenized *at3*^−/−^ background. We found that there was statistically significant survival in *at3*^−/−^;*f2*^*C/Δ*^ mutants, confirming that *frost* is a mutant allele of *f2* (Figure 3B).

**Figure 3.**
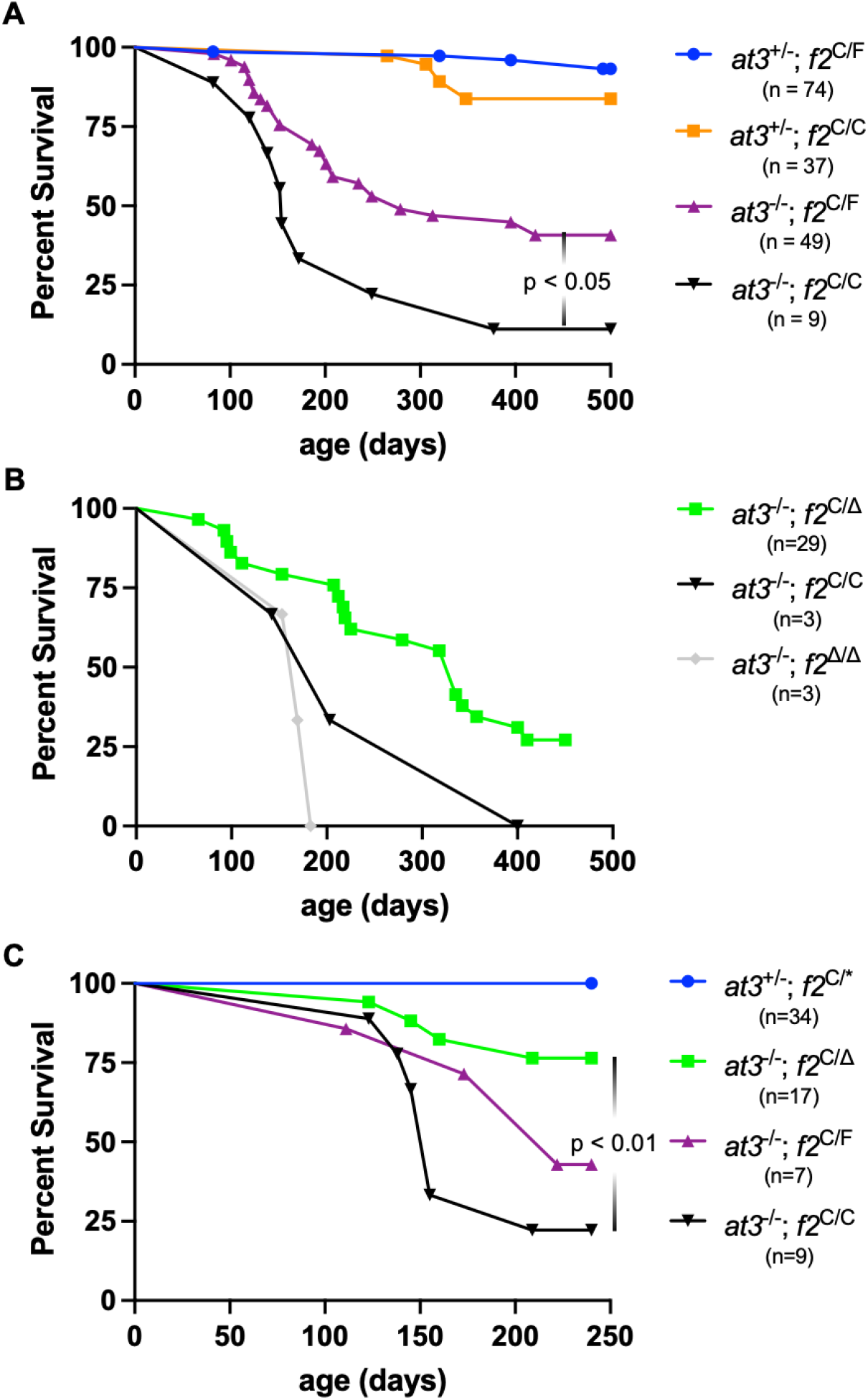
The *frost* allele is a loss-of-function variant in prothrombin that partially rescues survival due to At3 deficiency. (A) A single copy of *frost* (purple) significantly improves survival in *at3* deficient fish compared to wild-type (black) but does not return survival to the levels of heterozygous *at3* controls (blue and orange). (B) Observation of *f2*^*Δ*^ populations over time demonstrates that heterozygous fish survive longer than wild-type *at3* deficient fish, but that homozygosity for the modifier in each family results in increased mortality. (C) When compared in a single cohort of offspring, only the *f2*^*Δ*^ allele significantly rescues survival of the *at3* knockout compared to *f2*^*C/C*^. However, the *frost* allele does trend towards rescue and is not significantly different when directly compared to *f2*^*Δ*^. *f2*^*C/\**^ includes *f2*^*C/F*^ and *f2*^*C/Δ*^ genotypes.

A single cohort was generated to directly compare the ability of the *f2*^*Δ*^ and *frost* (*f2*^*F*^) alleles to rescue survival in *at3*-null offspring (Figure 3C). As expected, compound heterozygotes (*f2*^*Δ/F*^) had the lowest survival rates in both *at3*^+/−^ *and at3*^−/−^ backgrounds. Among *at3*^−/−^ fish, *f2*^*C/Δ*^ had the longer median survival compared to *f2*^*C/F*^. Notably, only *f2*^*Δ*^ and not the *frost* allele demonstrated a statistically significant survival benefit compared to the offspring with a prothrombin wild-type allele. However, when directly compared, the two alleles were not significantly different between each other. Overall, these genetic data suggest a difference in residual thrombin activity between *f2*^*Δ*^ and *f2*^*F*^, although in a limited cohort size.

### *frost* rescues the *at3* larval consumptive coagulopathy in a developmental-dependent manner

We have previously shown that loss of At3 results in a consumptive coagulopathy in the larval period^17^, and *at3*^−/−^ larvae are unable to form occlusive thrombi. To investigate the effects of *frost* on this phenotype, larvae were generated from an incross between *at3*^+/−^;*f2*^*C/F*^ parents and assayed using a laser-induced venous endothelial injury model at 3 dpf (Figure 4A). As expected, based on our prior *f2*^*Δ*^ mutant data^24^, homozygosity for *frost* (*f2*^*F/F*^) prevented occlusive thrombus formation independent of *at3* status. In *at3*^−/−^; *f2*^*C/C*^ larvae, clot formation was reduced as previously shown^17^. However, heterozygosity for *frost* (*f2*^*C/F*^) rescued the consumptive coagulopathy in 20% of fish. To confirm and further characterize the rescue, occlusive thrombus formation was analyzed at 3 time points (57, 78, and 101 hours post fertilization (hpf)) in *at3*^−/−^ larvae with differing *frost* genotypes (Figure 4B). At all three time points, roughly 10% of *f2*^*C/C*^ larvae and 0% of *f2*^*F/F*^ larvae demonstrated complete occlusion. In contrast, complete occlusion was seen in 13% of *f2*^*C/F*^ larvae at 57 hpf, and this rate increased to 24% at 83 hpf and 74% at 101 hpf. Overall, this suggests that a single copy of *frost* restores the ability to form clots in a developmentally-dependent manner in *at3*^−/−^ larvae due to a reduction in thrombinmediated fibrinogen consumption.

**Figure 4.**
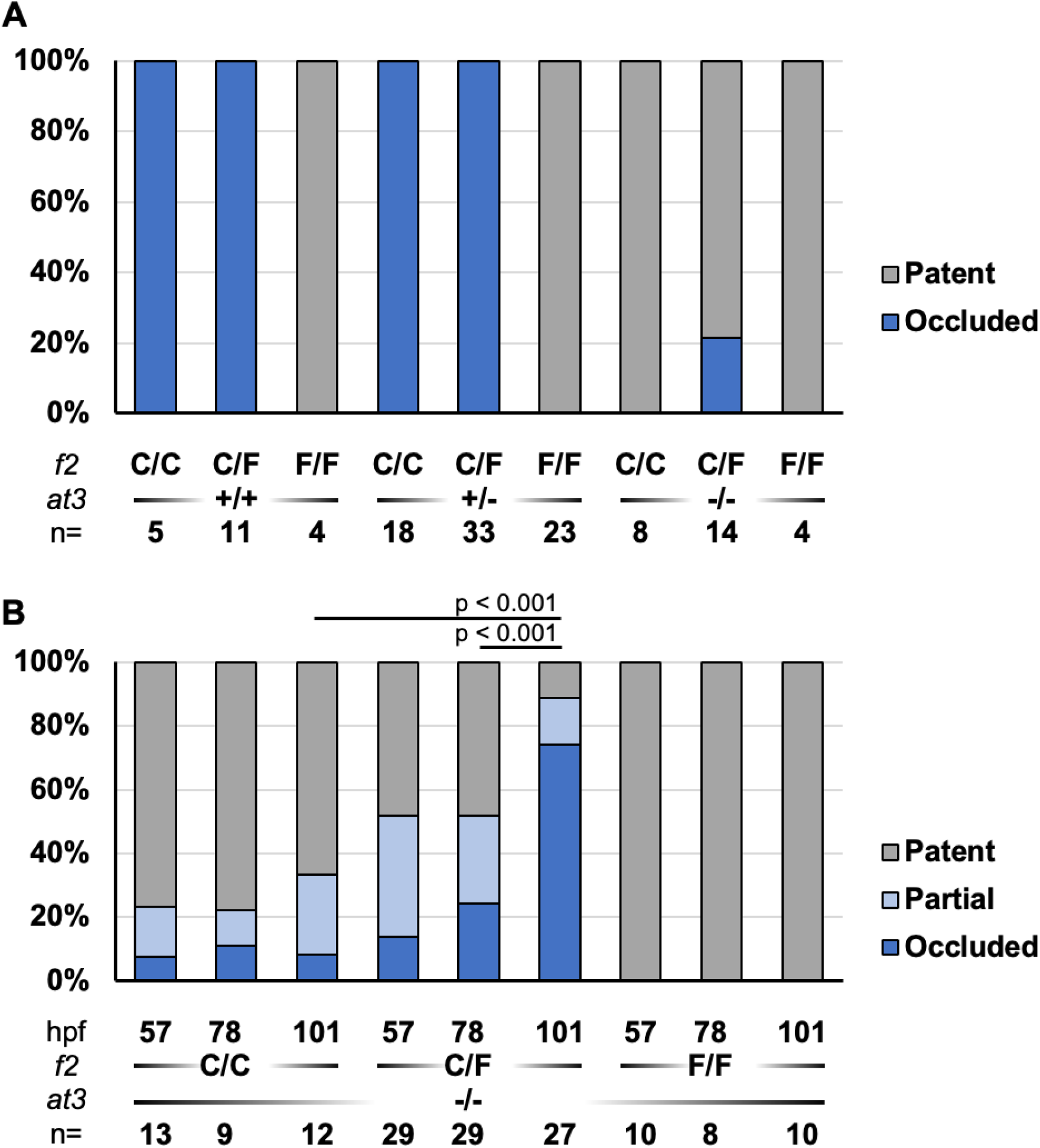
Heterozygosity for *frost* rescues the *at3* mutant consumptive coagulopathy in a developmental dependent manner. (A) Homozygosity for either the *at3* or *frost* mutations prevent vessel occlusion at 3 dpf following endothelial laser injury. Heterozygosity for the *frost* mutation partially rescues the ability of At3 deficient offspring to occlude. (B) A developmental time course of venous endothelial injury of offspring generated from an *at3*^−/−^*;f2*^*C/F*^ incross. Homozygosity for *frost* completely blocked thrombus formation while heterozygosity rescued the low occlusion rate of *at3* deficient larvae in a developmental dependent manner. Partial occlusion indicates the presence of vascular deposits or transient alterations in turbidity without the formation of an occlusive clot.

### Prothrombin secretion is reduced by the *frost* mutation

To interrogate how the *frost* mutation affects prothrombin activity, the homologous cysteine (C507) was mutated in a human prothrombin expression vector. As this mutation results in a free-thiol at C493, two additional prothrombin variants were generated; C493A and C493A/C507F. Expression of wild-type prothrombin in HEK293T cells resulted in secreted protein levels >100 ng/mL in the media. In contrast, all 3 cysteine mutants yielded levels <2 ng/mL (Figure 5A). In the corresponding cellular lysate, wild-type prothrombin transfected cells contained approximately 12 ng/mL prothrombin, while cells transfected with cysteine variants yielded levels 1.2-5.8 ng/mL (Figure 5B). This suggests that loss of the C493/C507 disulfide bond results in decreased secretion due to impaired protein production and/or increased degradation.

**Figure 5.**
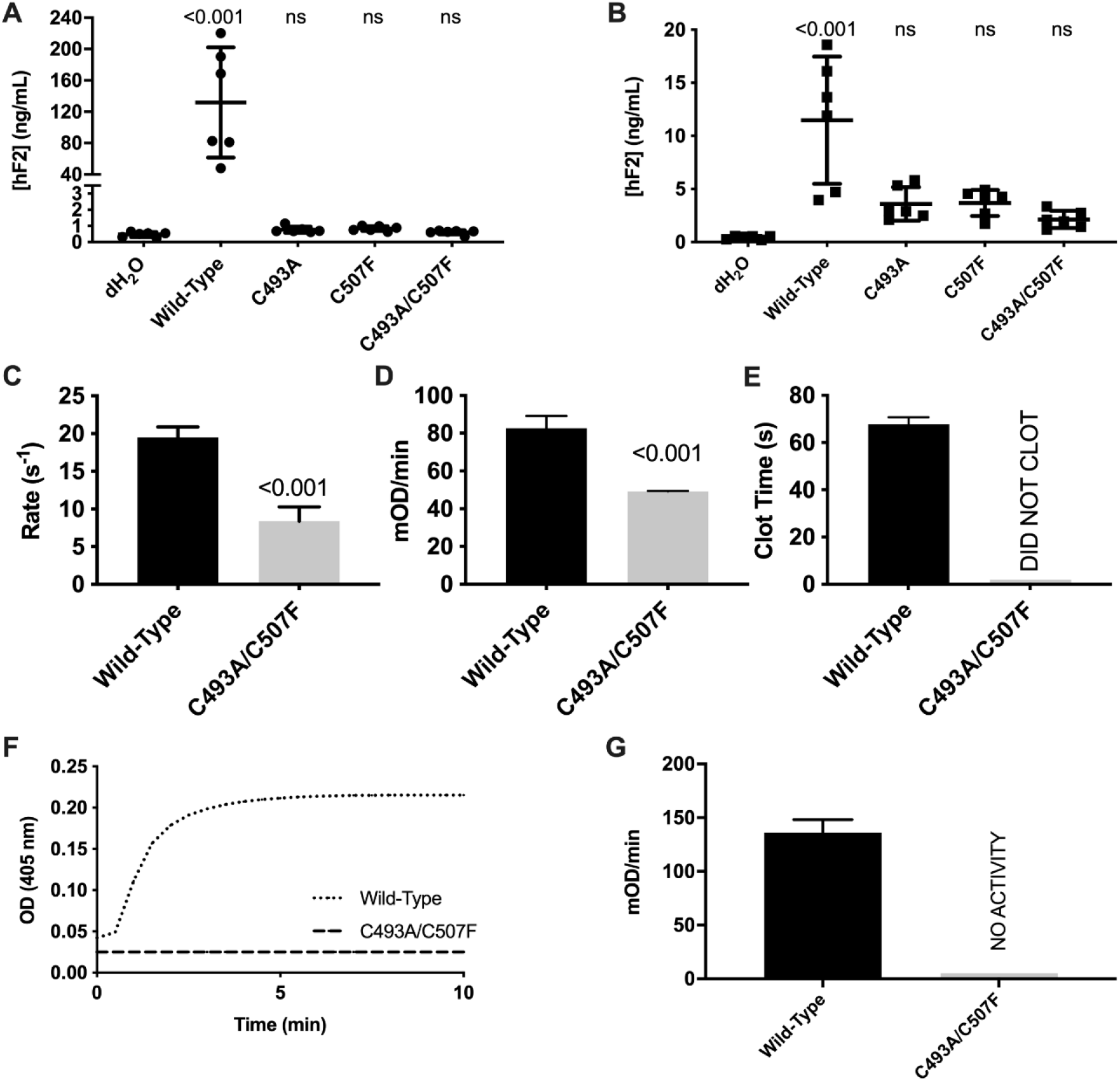
C507F mutation leads to reduced biosynthesis, activation, and activity of human prothrombin. The homologous mutation to *frost* was made in human prothrombin (C507F), expressed in HEK293T cells, and compared with and without a mutation in its paired cysteine at position 493 (C493A). Measurement by ELISA revealed protein levels indistinguishable from water controls in both the (A) media, and (B) cell lysate. (C) Thrombin generation by prothrombinase was significantly reduced when measured by DAPA fluorescence change. Fully activated prothrombin had reduced ability to (D) cleave the synthetic substrate S-2238, and (E) was unable to cleave fibrinogen. The rate was calculated during steady-state conditions and normalized to enzyme concentration. (F) Turbidity measurements showed no changes in the presence of mutant prothrombin. (G) The rate of clot formation could not be calculated for prothrombin C493A/C507F due to the inability to cleave fibrinogen.

### Loss of the C493/C507 disulfide bond results in impaired prothrombin activation and activity

The C493A/C507F prothrombin variant was purified from stably transfected HEK293 cells. Thrombin-DAPA complex formation demonstrated a decreased fluorescence signal change in mutant (1613 ± 91 RFU/µM) versus wild-type (3262 ± 347 RFU/µM), consistent with a disrupted protease domain. Additionally, the initial rate of activation was 2.2-fold slower in the mutant (8.4 ± 1.9 s^−1^) compared to the wild-type (19.5 ± 1.4 s^−1^) (Figure 5C).

The generated thrombin was assayed for proteolytic activity using S-2238 and fibrinogen as substrates. The amidolytic activity towards S-2238 was significantly decreased in the mutant thrombin (49.2 ± 0.2 mOD/min) compared to wild-type (82.7 ± 6.5 mOD/min), confirming a partially disrupted protease domain (Figure 5D).

Furthermore, C493A/C507F failed to cleave fibrinogen or alter the turbidity of the solution (Figure 5E-G). Overall, these data suggest a partially disrupted protease domain with reduced activity towards small peptidyl substrates and no activity towards macromolecular substrates like fibrinogen.

### Genetic insufficiency of additional coagulation factors mitigates thrombosis secondary to At3 deficiency

Complete genetic deficiency of *f10* in zebrafish has previously been shown to result in lethality by 3-4 months of age due to hemorrhage^22^. This line was bred separately to the At3-deficient strain to assess whether attenuation of other common pathway factors would modify thrombosis-induced lethality. Survival of these fish was tracked for 450 days, and in the *at3*^−/−^ background *f10*^+/−^ mutants demonstrated greater survival than the corresponding wild-type and homozygous mutant siblings (Figure 6A). This further supports the hypothesis that reductions in thrombin generation can improve survival in a prothrombotic background.

**Figure 6.**
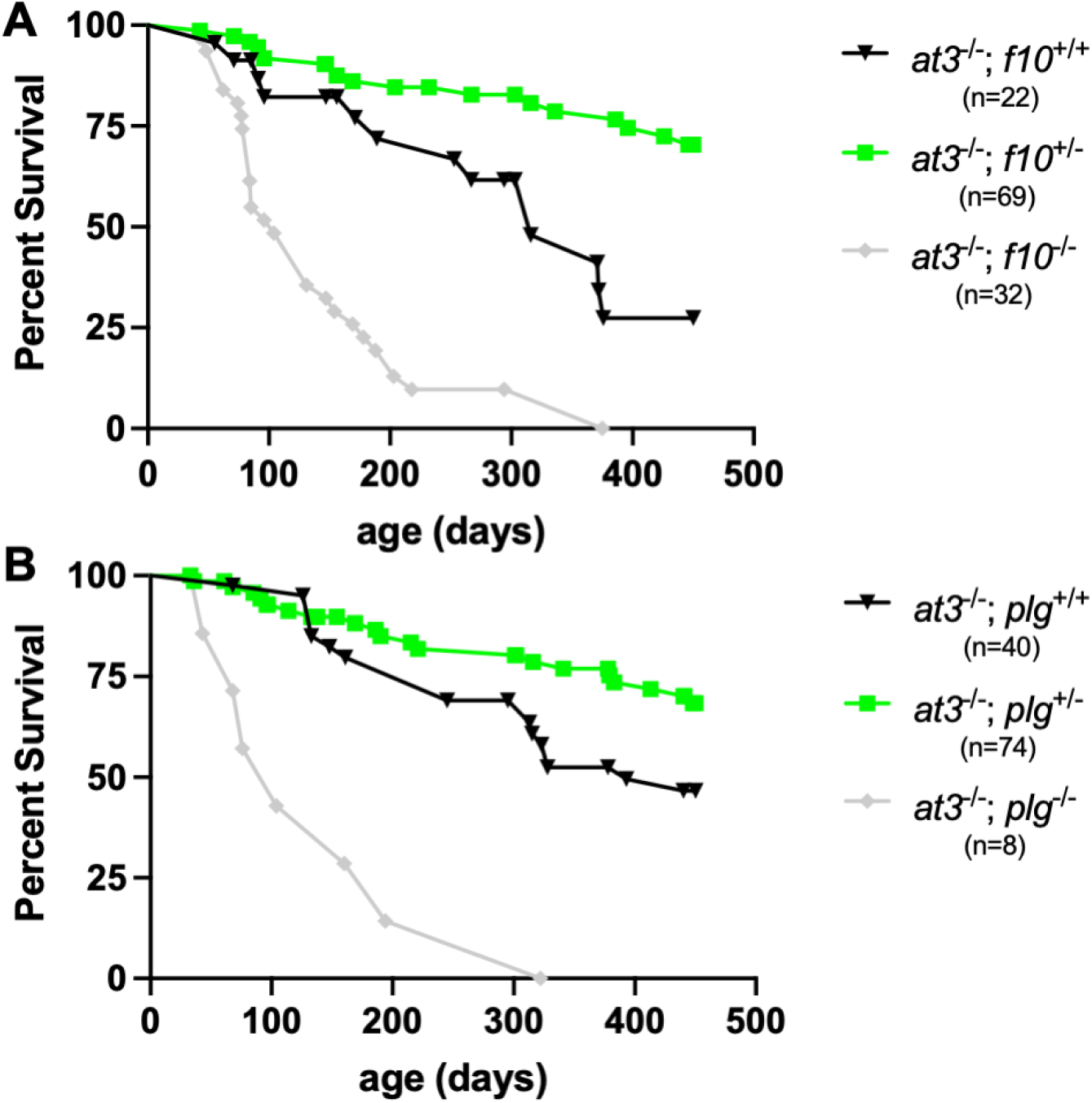
*f10 and plg* mutations protect against early lethality due to At3 deficiency. Observation of *f10* (A) and *plg* (B) populations over time demonstrates that heterozygous fish survive longer than wild-type *at3* deficient fish, but that homozygosity for the modifier in each family results in increased mortality.

Although *at3*^−/−^ adults are believed to succumb primarily due to thrombosis, there is a potential risk of bleeding due to the previously published hypofibrinogenemia^17^, therefore we evaluated the role of clot stability in this model. To interrogate fibrinolysis, *plg* mutants^23^ were mated into the *at3* mutant background. When tracked for survival, *at3*^−/−^*;plg*^+/−^ fish were found to have a modest advantage compared to *at3*^−/−^*;plg*^*+/+*^ fish (Figure 6B). Both live significantly longer than *at3*^−/−^*;plg*^−/−^ fish.

### Suppression of AT3-associated thrombosis by *F2* mutations was not detected in a large genotyped cohort

Using the UKBB cohort with its associated EHR (n=470,751: 7,992 VTE cases and 462,759 controls), we assessed VTE prevalence across mutually exclusive carrier groups for deleterious variants in *F2, AT3*, and *F10*. Sole carriers of the common *F2* G20210A mutation (G20210A^+^, n=361) showed a significantly higher prevalence of VTE compared with the large non-carrier reference group for *F2* and *AT3* (G20210A^−^, AT3del^−^, respectively, p=2.2×10^−16^) (Figure 7), underscoring the strong thrombotic risk associated with this well-known common mutation in individuals of European descent^4^. In contrast, sole carriers of deleterious variants in *AT3* (AT3del^+^, n=175, Figure 7) did not differ significantly from non-carriers.

**Figure 7.**
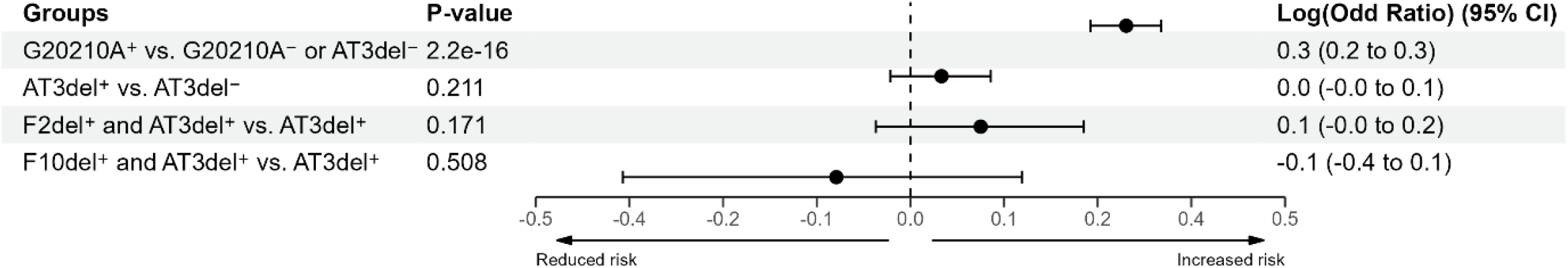
Association of deleterious coagulation factor variants with VTE in the UKBB. Individuals with HRC imputed genotype data (N=470,751) were classified as VTE cases (n=7,992) or controls (n=462,759) using ICD10 and procedure codes. Deleterious variants were identified using WGSA annotation and CADD score thresholds. Carrier groups were defined as follows: *F2* G20210A^+^ indicates carriers of the G20210A mutation; F2del^+^ indicates carriers of other deleterious *F2* variants (excluding G20210A); AT3del^+^ indicates carriers of *AT3* deleterious variants; and F10del^+^ indicates carriers of deleterious *F10* variants. Sole carriers were contrasted with corresponding non-carriers, and subgroup comparisons tested as to whether risk among AT3del^+^ individuals was modified by additional F2del^+^ or F10del^+^ mutations (see “Groups” column). VTE prevalence was compared using Fisher’s exact test, and results were visualized as a forest plot generated with the *forester* package in R.^41^

To evaluate potential effects of combined mutations, we compared AT3del^+^ carriers with and without additional variants in *F2* (excluding G20210A^+^) or *F10*. AT3del^+^ individuals carrying an additional *F2* deleterious mutation (AT3del^+^ and F2del^+^; n=68) or an additional *F10* mutation (AT3del^+^ and F10del^+^; n=13) were compared with those carrying *AT3* mutations alone. Neither subgroup analysis showed evidence that a second deleterious variant significantly altered VTE prevalence in AT3del^+^ individuals.

## Discussion

Although thrombosis is a leading cause of death worldwide, very few of the genetic risk factors are known. The increasing use of complex datasets to develop risk scores for thrombosis offers great potential but is limited to observational studies^12,34^. In this study we aimed to marry the discovery of new modifiers with the genetic accessibility of the zebrafish model to understand multigenic interactions.We have previously shown that complete loss of *at3* leads to early death by ~7 months of age in zebrafish^17^. We incorporated both forward and reverse genetic approaches to try to understand how hemostasis can be rebalanced in the context of thrombotic pathology. We performed an unbiased genome-wide chemical mutagenesis study. Having established four lines with survival in the context of complete At3 deficiency, we performed high throughput sequencing. Our initial candidate gene analysis using the Thrombogenomics consortium list uncovered an *f2* mutation at a conserved cysteine that we refer to as *frost*. This provided a proof-of-principle for the ability of the screen to identify modifier genes.

One of the most interesting findings of this study are the additional 3 lines identified, none of which carry a clearly identifiable deleterious mutation in a known clotting or clotting-related gene that segregates with survival. Based on our targeted knockouts, we would have expected to find additional suppressor lines due to mutations in *f10* or *plg*, since these also rescue the *at3*^−/−^ phenotype. We also should have found factor V gene mutations since their phenotypes virtually identical to the *f2* and *f10* mutants^18,22,24,35^. The fact that we did not observe any of these additional genes indicates that we did not achieve complete saturation of the genome and future mutagenesis experiments could identify additional new loci

Identification of the *frost* variant enabled mechanistic studies of thrombin in a C-terminal loop not previously studied. When looking specifically at mature C493A/C507F thrombin, the site of proteolysis is still maintained, albeit with reduced activity, as evidenced by S-2238 cleavage. In contrast, the ability to cleave fibrinogen is completely lost, suggesting a role in exosite-mediated recognition or stabilization of the interaction with macromolecular substrates. This may be explained by lower conformational stability around Y510 due to the lost disulfide bond resulting in decreased Na^+^ binding and indirectly altering exosite I^36,37^. Interestingly, when compared to our previous work^24^, the *frost* allele appears to have even less residual activity than our (*f2*^*Δ*^) targeted genetic knockout, consistent with the location of the conserved cysteine in the activated heavy chain.

When directly compared, the two *f2* alleles had differing abilities to rescue lethality in the *at3* knockout. Although our statistical interpretation was limited by cohort size, the survival trend suggests that the *frost* allele is not as potent of a suppressor. This is surprising given that *frost* shows less activity *in vitro* compared to the *f2*^*Δ*^ allele. Our previous study found that the *f2*^*Δ*^ allele maintained normal activity towards small-peptidyl substrates while having impaired but continued macromolecular activity (i.e. fibrinogen)^24^. This is not surprising given that the mutations from *f2*^*Δ*^ were imposed on the kringle 1 domain of prothrombin, which is absent from thrombin. In the case of *frost*, mutations of C493 and/or C507 lead to the loss of one of two intra B-chain disulfide bridges that span between D432 and S525 - two of the key catalytic triad residues of thrombin in prothrombin numbering. This led to a profound decrease in the proteolytic activity of thrombin whereby its activity towards small-peptidyl substrate was reduced by half while its procoagulant activity was completely abolished. Taken together, our data suggest that dampened thrombin activity due to heterozygous *f2* mutations were protective in the *at3* knockout, while complete loss of thrombin’s hemostatic potential is unable to rescue.

Having rescued survival with two independent *f2* alleles, we tested the effect on *at3*^−/−^ larval consumptive coagulopathy. While our endothelial-injury assay is normally performed at 3 dpf, initial studies yielded mixed results, with an incompletely penetrant rescue and an unpredictable inheritance pattern. Performing the assay at different times revealed an age-dependent rescue of the consumptive coagulopathy with increased penetrance of rescue at older ages. We have shown that in the *at3* knockout^17^, fibrinogen is consumed rapidly, presumably by basal uninhibited thrombin activity. However, in this study at subsequent timepoints, the continued production of fibrinogen seems to be able to overcome this consumption and re-establish hemostasis (Figure 4).

Using a reverse genetics approach, we knocked out two members of the common pathway, *f2* and *f10*^22,24^, that are endogenous targets of At3, and bred them into the *at3* mutant background. Consistent with the *frost* data, haploinsufficiency was able to partially rescue survival, but complete deficiency led to greater mortality independent of *at3* status. This is consistent with current clinical interventions that target thrombin and FXa for treatment of recurrent thrombosis^38^. We also explored the fibrinolysis system to understand the role of clot stability in survival. Surprisingly, heterozygosity of *plg* led to improved survival. This suggests that there may be an underappreciated risk of hemorrhage which might partially contribute to lethality in the *at3* knockout line and is ameliorated by *plg* deficiency. Indeed, we have observed sporadic hemorrhage in *at3* mutant adult fish^17^. This is consistent with reports supporting the use of a combination of the fibrinolysis inhibitor ε-aminocaproic acid and low dose heparin in the treatment of cases of disseminated intravascular coagulation with hyperfibrinolysis^39,40^. In addition, the median survival of wild-type controls in the *plg* survival study was around 1 year, which is longer than we have traditionally seen in the *at3* knockout line. This is likely due to the selection over time of background haplotypes that benefit *at3*^−/−^ survival in addition to variable environmental effects such as housing density and temperature that can greatly affect zebrafish growth. This concern is mitigated by our use of sibling controls, which control for such variability.

The finding that variation in prothrombin or FX modify antithrombin deficiency was not surprising, and in fact predictable. Therefore, we expected to observe a similar phenomenon in a large-scale, epidemiological resource like the UKBB. We saw the very strong expected association of thrombosis with the common *F2* G20210A variant. However, we did not observe significantly different rates of VTE in *AT3* variants, and consequently we could not identify the modifying effect of *F2* or *F10* variation. The most likely explanation is that even with this large database, the low frequency of AT3 deficiency resulted in it being underpowered to detect associated thrombosis. This demonstrates that even a large human dataset does not have adequate power to study these protein coding mutations. We were only able to observe this very logical and expected association in the zebrafish model where the power to detect such an association is notably higher. This does not suffer from survival or selection biases decreasing power, which are common in human Biobank based datasets.

Our work continues to demonstrate the use of zebrafish in studying human thrombosis. Not only were we able to prove the conservation of re-balancing hemostasis to mitigate thrombosis through *f2* and *f10* mutation, but we were able to confirm an unexpected role of fibrinolysis as well. The *frost* mutation is proof that a genome-wide suppressor experiment is able to identify modifiers of thrombosis, and the other 3 lines provide strong support for the existence of non-canonical modifying factors. *frost* also indicates that even in the era of massive human databases, high-throughput genetic vertebrate animal models like the zebrafish are still needed for discovery of low frequency modifying effects. Once we are able to map and clone the suppressor mutations in the additional lines, we expect that they will identify novel factors that regulate coagulation. Further work to map novel loci offers great potential in advancing our understanding of coagulation, with implications for diagnostics and therapeutics for people at risk of and affected by thrombosis.

## Acknowledgements

The authors thank the University of Michigan Advanced Genomics Core for services. UKBB data research has been conducted using the UK Biobank Resource under application number 24460.

## Author contributions

S.J.G. designed and performed research, analyzed data and wrote the manuscript; Y.L. and A.H.V. designed and performed research, and analyzed data. C.E.R., X.Y., A.R.D., A.C.F., and H.M. performed research. M.Y., I.S., P.Y.K. and C.A.K. designed research, analyzed data, and edited the manuscript; J.A.S. designed, performed, and supervised research, analyzed data, and wrote the manuscript. All authors reviewed the manuscript.

